# A multicenter, randomized study of decitabine as epigenetic priming with induction chemotherapy in children with AML

**DOI:** 10.1101/124263

**Authors:** Lia Gore, Timothy J. Triche, Jason E. Farrar, Daniel Wai, Christophe Legendre, Gerald C Gooden, Winnie S. Liang, John Carpten, David Lee, Frank Alvaro, Margaret E. Macy, Philip Barnette, Todd Cooper, Laura Martin, Aru Narendran, Jessica Pollard, Soheil Meshinchi, Jessica Boklan, Robert J Arceci, Bodour Salhia

## Abstract

**Background:** Decitabine is a deoxycytidine nucleoside derivative inhibitor of DNA-methyltransferases, which has been studied extensively and is approved for myelodysplastic syndrome in adults but with less focus in children. Accordingly, we conducted a phase 1 multicenter, randomized, open-label study to evaluate decitabine pre-treatment before standard induction therapy in children with newly diagnosed AML to assess safety and tolerability and explore a number of biologic endpoints.

**Results:** Twenty-four patients were fully assessable for all study objectives per protocol (10 in Arm A, 14 in Arm B). All patients experienced neutropenia and thrombocytopenia. The most common grade 3 and 4 non-hematologic adverse events observed were gastrointestinal toxicities and hypophosphatemia. Plasma decitabine PK were similar to previously reported adult data. Overall CR/CRi was similar for the two arms. MRD negativity at end-induction was 85% in Arm A versus 67% in Arm B patients. DNA methylation measured in peripheral blood over the course of treatment tracked with blast clearance and matched marrow aspirates at day 0 and day 21. Unlike end-point marrow analyses, promoter methylation in blood identified an apparent reversal of response in the lone treatment failure, one week prior to the patient’s marrow aspirate confirming non-response. Decitabine-induced effects of end-induction marrows in Arm A were reflected by changes in DNA methylation and gene expression comparison with matched paired marrow diagnostic aspirates.

**Conclusions:** This first-in-pediatrics trial demonstrates that decitabine prior to standard combination chemotherapy is feasible and well tolerated in children with newly diagnosed AML. Pre-treatment with decitabine may represent a newer therapeutic option for pediatric AML, especially as it appears to induce important epigenetic alterations. The novel biological correlates studied in this trial offer a clinically relevant window into disease progression and remission. Additional studies are needed to definitively assess whether decitabine can enhance durability responses in children with AML. This trial was registered at www.clinicaltrials.gov as NCT01177540.

## BACKGROUND

Attaining complete response/remission (CR) is currently considered the essential first step in the effective treatment of acute myelogenous leukemia (AML). Historically, the most widely used induction therapy included seven days of cytarabine plus three days of anthracycline (known as “7+3”). With this approach, 75-80% of children with AML achieve CR (1-3). Subsequently, the addition of a third agent such as etoposide to 7+3 (ADE) along with expanded supportive care measures, has led to higher remission induction rates of approximately 85%. Of patients who do not attain remission, approximately one-half have resistant leukemia and a substantial proportion will die from complications of the disease or treatment. Thus, there is a need to develop new treatment strategies to improve outcomes for these patients.

Pediatric tumors have been shown to have lower mutation burdens than adult tumors and many of these mutations occur in the plethora of known epigenetic complexes (4). In addition, significant aberrant DNA methylation is also observed in pediatric cancers such as AML including in patients with the poorest risk sub-types (5). These studies argue for the importance of identifying novel epigenetic therapies that target both histone and/or DNA methylation modifications. Specifically, reversal of promoter DNA hypermethylation and associated gene silencing is an attractive therapeutic approach in adult cancers. The DNA methylation inhibitors decitabine and azacitidine are efficacious for hematological neoplasms at lower, less toxic, doses (6). Experimentally, high doses induce rapid DNA damage and cytotoxicity, which do not explain the prolonged response observed in adult patients (6). Studies have consistently shown that transient low doses of DNA demethylating agents exert durable anti-tumor effects on hematological and epithelial tumor cells (6). Moreover, studies have demonstrated that DNA hypomethylating agents can sensitize resistant cancer cells to cytotoxic agents *in vitro* and *in vivo (7-15)* and can enhance chemosensitivity of human leukemia cells to cytarabine (16). Therefore, pretreatment with a DNA hypomethylating agent may increase the efficacy of pediatric AML induction therapy (17). However, to date there are no studies to demonstrate the safety, tolerability, or any signal of efficacy of decitabine in combination with conventional multi-agent chemotherapy for AML in children. We report here the first phase 1 clinical evaluation of decitabine in children with newly diagnosed AML as a feasibility study to determine the safety, tolerability, and preliminary efficacy when used as epigenetic pretreatment before induction chemotherapy. In addition to assessing toxicity and morphologic remission, this study examined decitabine pharmacokinetics and minimal residual disease (MRD) impact. We also performed global DNA methylation and RNA-seq analyses to examine how decitabine priming impacted the methylome and gene expression in end-induction marrows when compared with matched diagnostic marrow baseline controls. We believe that this feasibility study was essential prior to longer-term studies assessing whether epigenetic-directed therapy in pediatric AML can lead to enhanced response rates or more durable responses.

## PATIENTS, MATERIALS, AND METHODS

### Patient eligibility

Eligible patients were 1 to 16 years of age (inclusive), had histologically confirmed *de novo* AML with >20% bone marrow blasts and adequate cardiac function (defined as ejection fraction >50% or shortening fraction >26%). Patients with acute promyelocytic leukemia (FAB M3 subtype), symptomatic CNS involvement, white blood cell count over 100,000/μl, significant renal or hepatic disease, any prior chemotherapy or radiation therapy for AML, known HIV infection, history of CML, congenital syndromes known to predispose to AML (for example, Down syndrome, Fanconi anemia, Kostmann syndrome, or Diamond-Blackfan anemia) were excluded.

The study protocol was approved by the institutional review boards at participating sites and was conducted in accordance with the Declaration of Helsinki, Good Clinical Practice, and all local and federal regulatory guidelines. A parent or legal guardian provided written informed consent, with patient assent as appropriate according to institutional requirements.

### Study design

This multicenter, open-label study randomized patients to one of two arms: either five days of decitabine followed by standard induction chemotherapy with cytarabine, daunorubicin, and etoposide (Arm A=DADE), or standard induction chemotherapy with cytarabine, daunorubicin, and etoposide without decitabine (Arm B=ADE). The trial was listed under ClinicalTrials.gov identifier: NCT00943553. Twenty-five children age 1-16 years with newly diagnosed *de novo* AML were randomized to receive either Arm A or Arm B. Given the feasibility nature of the study, sample size was selected based on the likelihood of how many patients might be accrued in a reasonable time frame so that future studies could be planned. Patients were stratified by age group and then randomized within each stratum in a 1:1 ratio by an Interactive Voice Response System via a random number generator. Three age strata were used: 1 to <2 years, 2-11 years, and 12-16 years, with efforts made to balance enrollment among the age groups.

All patients received one cycle of study treatment in the absence of clinically significant disease progression, unacceptable toxicity, or patient/guardian choice to discontinue participation. Patients were not pre-medicated prior to the first dose of decitabine; however, all other supportive care measures were allowed according to institutional standards. Following the completion of the study therapy, therapy continued at the treating physician’s discretion.

Treatment was administered to patients in hospital, and hospitalization through count recovery was mandated. The dose and schedule of decitabine used in this study was known to be safe and tolerable in adults and was known to induce adequate hypomethylation (18, 19), inhibit DNA methyltransferase, and induce tumor suppressor gene activation as early as 3-5 days following initiation. Treatment included: a) decitabine 20 mg/m^2^ IV infusion for 1 hour daily for 5 days (Arm A) on Days 1-5; b) age-based dosing of intrathecal cytarabine (1 to <2 years: 30 mg; 2 to <3 years: 50 mg; ≥3 years: 70 mg) at the time of diagnostic lumbar puncture or on Day 1; c) cytarabine 100 mg/m^2^/dose (3.3 mg/kg/dose for BSA <0.6 m^2^) slow IV push over 15 minutes, every 12 hours for 10 days on Days 1-10 (Arm B) or Days 6 to 15 (Arm A); d) daunorubicin 50 mg/m^2^ (1.67 mg/kg/dose for BSA <0.6 m^2^) IV over 6 hours for 3 days on Days 1, 3, 5 (Arm B) or Days 6, 8 and 10 (Arm A); and e) etoposide 100 mg/m^2^/dose (3.3 mg/kg/dose for BSA <0.6 m^2^) IV over 4 hours for 5 days on Days 1-5 (Arm B) or Days 6-10 (Arm A).

Toxicity was graded according to the National Cancer Institute Common Terminology Criteria for Adverse Events (CTCAE), version 4.0 (http://ctep.cancer.gov; National Cancer Institute, Bethesda, MD). Treatment-related toxicity was defined as non-resolving grade 3 or grade 4 non-hematologic or hematologic toxicity or time to platelet recovery to ≥100,000/μl and neutrophil recovery to ≥1,000/μl more than 55 days from the last day of induction chemotherapy in the absence of leukemia. Events considered by the investigator to be possibly, probably, or definitely related to decitabine were considered treatment-related toxicity.

### Safety assessments

Induction mortality was defined as death occurring within six weeks following initial diagnosis of AML. An independent Data Safety and Monitoring Board assessed the first 12 patients enrolled. This Board remained active for continuous analyses and recommendations throughout the conduct of the study. Stopping rules were included in the protocol to ensure appropriate safety of participants and that in the event of unacceptable toxicity additional patients would not be placed at risk. All investigators had access to the primary clinical trial data.

### On-study evaluations

Required assessments included physical examinations and recording of adverse events at screening/baseline, on day 5, and at the completion of study therapy. Required hematology and serum chemistry assessments were performed on days 1, 2, 6, 7, 14, 15, and weekly thereafter. Bone marrow evaluations for morphology, MRD and molecular analyses were performed at screening/baseline, 3-4 weeks following the completion of induction chemotherapy regardless of peripheral blood count recovery, and then as clinically indicated until count recovery. Any clinically appropriate assessment or test was allowed at the treating physician’s discretion to maintain standards of care.

### Efficacy assessments

The primary efficacy variable was Complete Response (CR), defined by the International Working Group 2003 criteria (20), requiring patients to have a morphologic leukemia-free state and an absolute neutrophil count >1000/μL and platelets of > 100,000/μL. Neither hemoglobin nor hematocrit was considered to have bearing on response although patients were required to be red blood cell transfusion independent to enroll. Secondary efficacy variables included Leukemia-Free Survival (LFS), Overall Survival (OS), methylation of DNA following decitabine therapy, times to platelet and neutrophil recovery, and level of minimal residual disease at the end of induction therapy. LFS and OS were assessed on patients every three months until disease progression, death, or loss to follow up. MRD analysis was performed at the post-induction therapy assessment by multi-parameter flow immunophenotyping. Due to the small sample size, statistical analyses were primarily descriptive.

### Pharmacokinetic evaluations

Serial blood samples (2 mL each) were drawn from all patients randomized to Arm A at pre-decitabine, 30, 60 (just prior to end of infusion), 65, 90, 120, and 180 minutes after the start of decitabine infusion. A separate line was used to draw PK samples not in proximity (i.e., not the contralateral lumen of a double lumen line) to the decitabine infusion. Samples were collected in EDTA tubes containing tetrahydrouridine, a cytidine deaminase inhibitor, to prevent decitabine degradation, and were centrifuged at 4C within 30 minutes of collection. Plasma was harvested and stored frozen at -70 to -80C and shipped on dry ice for central analysis.

Pharmacokinetic parameters were calculated from plasma decitabine concentration-time data by non-compartmental methods using Phoenix WinNonlin version 6.2 (Pharsight Corporation, Mountain View, CA). The maximum plasma concentration (C_max_) and the time at which C_max_ occurred (T_max_) were determined by inspection of the individual data. AUC from time 0 until the last quantifiable concentration (AUC_0–tau_) was determined by the linear up-log down trapezoidal rule. The terminal phase elimination rate constant (K_el_) was estimated from the slope of the concentration-time data during the log-linear terminal phase using least square regression analysis. The terminal phase elimination half-life (t_1/2_) was calculated using the formula 0.693/Kel. The AUC-time curve from 0 to infinity (AUC_0-infinity_) was computed as AUC_0-t_ plus the extrapolation from the last quantifiable concentration, C_t_, to infinity using the formula C_t_/K_el_. Total body clearance (CLp) was calculated by the formula Dose/AUC_0-infinity_. The volume of distribution at steady state (V_dss_) was calculated using the formula CL_p_ x MRT. Area under the first moment curve (AUMC) was determined using the linear trapezoidal rule to calculate AUMC_0-tau_ and extrapolated to infinity as AUMC_0-tau_ + t * C_t_/(K_el_)^2^. The formula used to determine mean residence time (MRT) was [AUMC/AUC_0-infinity_–tau/2], where tau is the duration of infusion.

### DNA methylation analysis

Bone marrow and blood samples were obtained from all patients at baseline and at completion of induction therapy. In addition, blood samples were also collected on days 7 and 14. DNA was extracted from marrow or peripheral blood lymphocytes (buffy coat) using Qiagen’s AllPrep kit from samples enriched for leukemic blasts by standard Ficoll separation. Global DNA methylation was evaluated using the Infinium® Human Methylation450® BeadChip Array according to the manufacturer’s protocol (Illumina, San Diego, CA) and as previously described (21). A total of 18 paired patient samples with both diagnostic and remission bone marrows (9 pairs from Arm A and 9 pairs from Arm B, totaling 36 samples) were used for DNA methylation analyses. In addition, peripheral blood DNA from all time points was also analyzed. DNA Methylation levels for each CpG residue are presented as β values, estimating the ratio of the methylated signal intensity over the sum of the methylated and unmethylated intensities at each locus. The average β value reports a methylation signal ranging from 0 to 1 representing completely unmethylated to completely methylated values, respectively. DNA methylation data were preprocessed using the Illumina Methylation Analyzer (IMA; doi: 10· 1093/bioinformatics/bts013), including background and probe design corrections, quantile normalization and logit transformation. Loci with detection p-values >0.05 in 25% of samples, on sex chromosomes, or within 10 bp of putative SNPs were removed from analysis. Differential methylation analysis was performed by IMA. A paired Wilcoxon rank test was conducted to compare end-induction marrows with diagnostic marrows within each arm. Probes with *p* < 0.05 having group-wise differences in β values of at least 0.15 were considered statistically significant and differentially methylated. Differentially methylated loci were visualized on a heat map and separation of groups was assessed by hierarchical cluster analysis using Manhattan distance and Ward’s method. Unsupervised clustering was also performed on the top 0.1% most variable probes by standard deviation. The DNA methylation data discussed here were deposited in NCBI Gene Expression Omnibus Database and are accessible through GEO Series accession number GSE78963.

### RNA sequencing analysis

Total RNA was isolated using Qiagen’s AllPrep kit and used to generate whole transcriptome libraries using Nugen’s Ovation RNA-Seq System v2 and KAPA Biosystems’ Library Preparation Kit with Illumina adapters. Equimolar libraries were pooled and used to generate clusters on Illumina HiSeq Paired End v3 flowcells on the Illumina cBot using Illumina’s TruSeq PE Cluster Kit v3. Clustered flowcells were sequenced by synthesis on the Illumina HiSeq 2000 using paired-end technology and Illumina’s TruSeq SBS Kit, extending to 83 bp for each of two reads and a 7bp index read. Approximately 75-100M reads per library were targeted for generation. Following sequencing, BCL files were converted to FASTQs using Illumina BCLConverter tool. FASTQ alignments were performed using Star to generate individual BAM files. Cufflinks/Cuffdiff and DEseq2 were used to identify differentially expressed transcripts. Annotations were based on Gencode version 3 (ENSEMBL) build 37.1. The RNA-seq data discussed in this publication have been deposited in NCBI’s Gene Expression Omnibus, and are also accessible through the GEO series accession GSE78963

### Analysis of repeat element transcription

Kallisto (22) was used to quantify abundance of transcripts using ENSEMBL build 80 and human repeat exemplars from RepBase 20_07 (23). Transcripts were quantified in units of transcripts per million (TPM). The resulting estimates were imported using the Artemis package (24) and annotated for presentation using TxDbLite (25), the latter providing a “best fit” alignment of RepeatMasker and RepBase element-wise repeat phylogeny.

### Pathway analysis

Gene lists of interest were uploaded into IPA (Ingenuity® Systems, Redwood City, CA) and the Core Analysis workflow was run with default parameters. Core Analysis provides an assessment of significantly altered pathways, molecular networks and biological processes represented in the samples’ gene list.

## RESULTS

### Patients

Twenty-five patients, aged 1-16 years (median 8.0 years) with WBC at diagnosis ranging from 1.19-58.09 x 10^3^/μL were randomized between March and November 2011. Two patients did not receive the full induction regimen due to toxicity. As shown in **Table 1**, 24 were fully assessable for all study objectives per protocol (10 in Arm A, 14 in Arm B). Three patients had confirmed FLT3 internal tandem duplications, all with an allelic ratio of ≥0.5 and one had a FLT3 D835 point mutation. Two patients had NPM1 mutations and two patients had CEBPA mutations. No patients had mutations of TET2, IDH1, IDH2, or C-CBL exons 8 or 9. One patient each had a KIT exon 8 (N822K) and 17 (D816H) mutation. Three patients had WT1 exon 7 mutations and one patient had a WT1 exon 9 mutation. On relative dose intensity analysis, patients received 99-100% of the intended doses of decitabine, daunorubicin, and etoposide and 84% of the intended doses of cytarabine.

**Table 1.** Patient Characteristics (by arm and overall)

| Characteristic | Patients | ...in Arm A | ...in Arm B |
| --- | --- | --- | --- |
| Consented | 25 (100%) | 11 (44%) | 14 (56%) |
| Randomized and treated | 25 (100%) | 11 (44%) | 14 (56%) |
| Evaluable for all study endpoints | 24 (96%) | 10 (40%) | 14 (56%) |
| Sex: |  |  |  |
| Male | 12 (48%) | 7 (58%) | 5 (42%) |
| Female | 13 (52%) | 4 (31%) | 9 (69%) |
| Age, years |  |  |  |
| Median (range) | 8.0 (1-16) | 8 (2-16) | 7.5 (1-16) |
| Race: |  |  |  |
| White | 20 (80%) | 8 (32%) | 12 (48%) |
| African-American | 1 (4%) | 1 (4%) | 0 (0%) |
| Asian | 1 (4%) | 0 (0%) | 1 (4%) |
| American Indian or Alaska native | 1 (4%) | 1 (4%) | 0 (0%) |
| Other | 2 (8%) | 1 (4%) | 1 (4%) |
| Ethnicity, Hispanic | 6 (24.0) |  |  |
| Ethnicity, not Hispanic or Latino | 19 (76.0) |  |  |
| Cytogenetic Risk Groups: |  |  |  |
| Favorable | 4 (8%) | 1 (4%) | 3 (12%) |
| Intermediate | 17 (68%) | 6 (24%) | 11 (44%) |
| Unfavorable | 4 (16%) | 4 (16%) | 0 (0%) |
| Cytogenetics: |  |  |  |
| Normal | 7 (28%) | 3 (12%) | 4 (16%) |
| t(8;21) | 4 (16%) | 1 (4%) | 3 (12%) |
| 11q23 | 9 (36%) | 5 (20%) | 4 (16%) |
| Other | 5 (20%) | 2 (8%) | 3 (12%) |
| Location: |  |  |  |
| Alberta (Canada) | 1 (4%) | 1 (4%) | 0 (0%) |
| Children's Hospital Central California | 1 (4%) | 0 (0%) | 1 (4%) |
| Denver | 8 (32%) | 5 (20%) | 3 (12%) |
| Emory | 1 (4%) | 0 (0%) | 1 (4%) |
| Johns Hopkins | 5 (20%) | 5 (20%) | 0 (0%) |
| John Hunter Hospital (Australia) | 2 (8%) | 0 (0%) | 2 (8%) |
| Mayo Clinic | 1 (4%) | 1 (4%) | 0 (0%) |
| Nationwide Children's Hospital | 1 (4%) | 0 (0%) | 1 (4%) |
| Phoenix Children's Hospital | 1 (4%) | 0 (0%) | 1 (4%) |
| Primary Children's Hospital | 1 (4%) | 0 (0%) | 1 (4%) |
| Royal Children's Hospital (Australia) | 3 (12%) | 1 (4%) | 2 (8%) |

### Toxicity

Treatment-emergent AEs are summarized in **Table 2**. The most common grade 3 and grade 4 AEs were hematologic, including WBC decreased, anemia, platelet count and neutrophil count decreased. Colitis (n=2), anorexia (n=3), hypophosphatemia (n=2), and hypokalemia (n=3) were also noted. One patient in Arm A experienced colonic perforation on Day 6 due to leukemic infiltration of the bowel wall that led to study discontinuation. Two patients in Arm A died 6 months after completion of induction therapy; one of necrotic bowel and *Pseudomonas* sepsis, and one of multisystem organ failure. The latter patient died 5 months after study treatment as a complication of stem cell transplantation. Neither death was attributed to decitabine nor to the chemotherapy regimen received during study participation.

**Table 2.** Grade 3 and Grade 4 treatment-emergent adverse events (TEAEs) reported in treated patients fully assessable for all study endpoints, as assessed by the Common Terminology Criteria for Adverse Events, version 4.0

|  | Decitabine + Chemotherapy |  | Chemotherapy only |  |
| --- | --- | --- | --- | --- |
| Adverse event | Grade 3<br>n (%) | Grade 4<br>n (%) | Grade 3<br>n (%) | Grade 4<br>n (%) |
| Any Grade 3 or Grade 4 TEAEs | 7 (87.5) | 7 (87.5) | 7 (77.8) | 7 (77.8) |
| **Blood and lymphatic system disorders** |  |  |  |  |
| Anemia | 6 (75.0) | 2 (25.0) | 4 (44.4) | 1 (11.1) |
| Febrile neutropenia | 1 (12.5) | 0 | 4 (44.4) | 0 |
| Other: Neutropenia | 1 (12.3) | 3 (37.5) | 1 (11.1) | 1 (11.1) |
| Other: Thrombocytopenia | 4 (50.0) | 4 (50.0) | 0 | 2 (22.2) |
| **Gastrointestinal disorders** |  |  |  |  |
| Colitis | 2 (25.0) | 0 | 2 (25.0) | 0 |
| **Investigations** |  |  |  |  |
| Neutrophil count decreased | 0 | 0 | 1 (11.1) | 3 (33.3) |
| Platelet count decreased | 0 | 1 (12.5) | 2 (22.2) | 3 (33.3) |
| White blood cell count decreased | 6 (75.0) | 6 (75.0) | 3 (33.3) | 5 (55.6) |
| **Metabolism and nutrition disorders** |  |  |  |  |
| Decreased appetite | 3 (37.5) | 0 | 0 | 0 |
| Hypokalemia | 3 (37.5) | 0 | 1 (11.1) | 0 |
| Hypophosphatemia | 2 (25.0) | 0 | 0 | 0 |

### Pharmacokinetics

Plasma concentrations of decitabine were quantifiable in all patients up to the last time point of 180 minutes. Post-infusion, plasma concentrations declined in a bi-exponential manner (**Figure 1**). Selected PK parameters of decitabine in subjects overall are shown in **Table 3**. The overall mean (standard deviation) PK parameters for the decitabine-treated patients were: C_max_, 294(104) ng/mL; AUC_0-∞_, 214(72.4) ng•h/mL; CL, 128(92.3) L/h; Vd_ss_, 45.5(41.1) L; 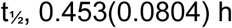; t_max_, 0.831h(0.253). Estimated PK values for a 70-kg adult male receiving a 1 hour decitabine 20 mg/m^2^ infusion are shown for reference. The mean exposure to decitabine, as measured by C_max_ and AUC was similar in patients aged 12-16 years compared with those aged 2-11 years as shown, and similar to those in adults. However, inter-patient variability in this study was high.

**Figure 1.**
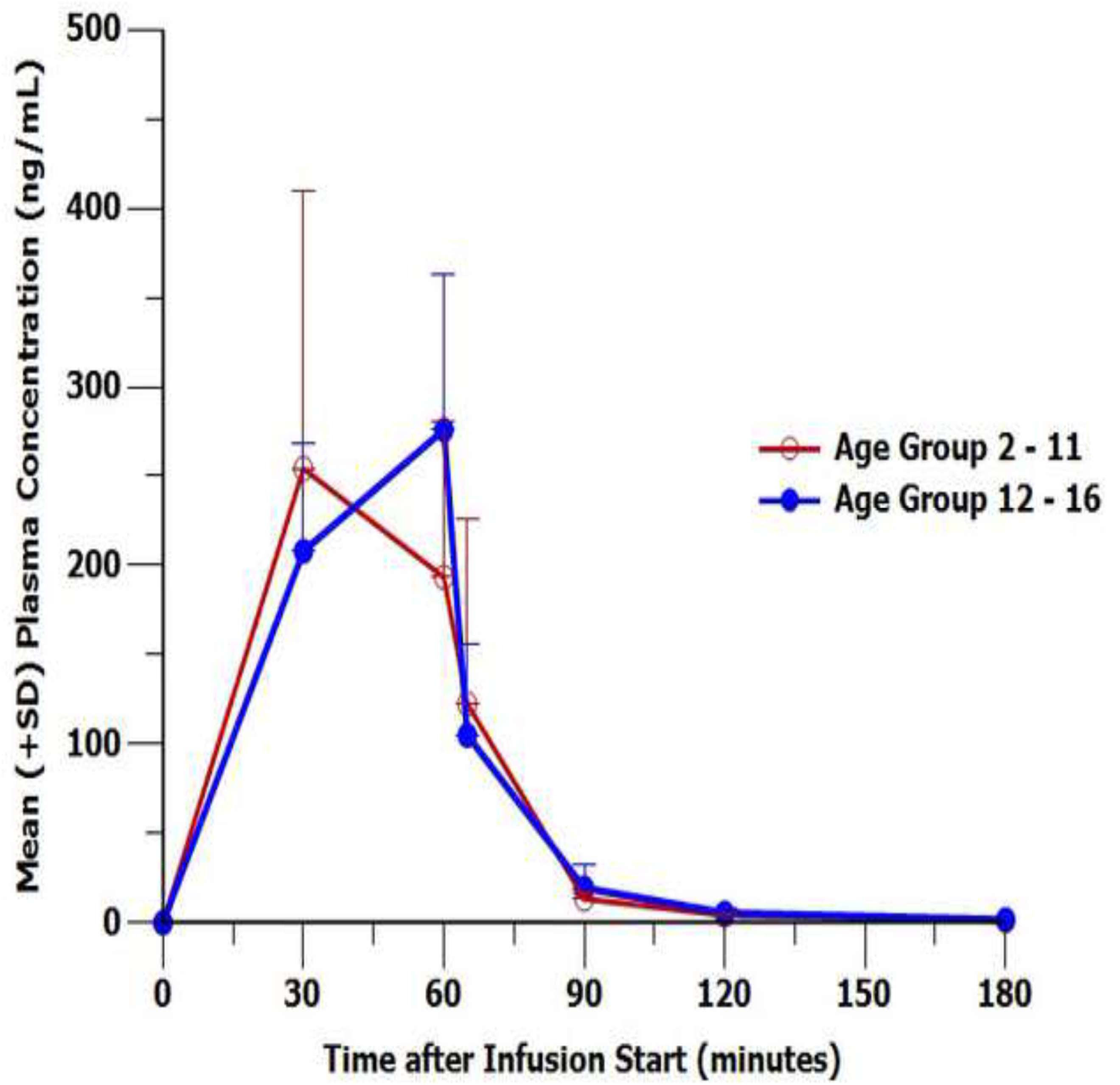
Mean decitabine blood concentration-time profile measured in whole blood by age group.

**Table 3.**
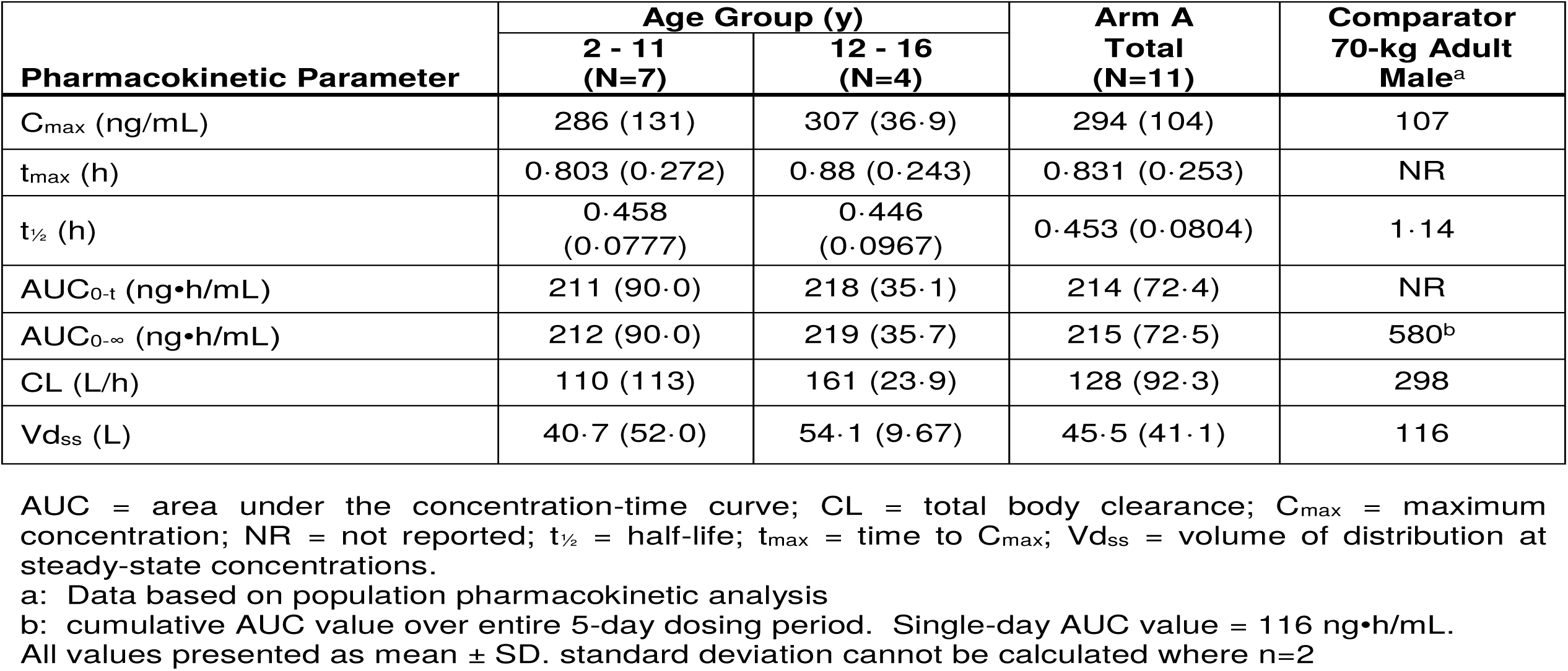
Mean Pharmacokinetic Parameters of Decitabine on Day 5 of Treatment—Overall and by Age Group, Pharmacokinetic Analysis Population

### Anti-leukemic response

Morphologic CRs and CRs with incomplete count recovery (CRi) rates were similar in both treatment arms: 100% CR/CRi in Arm A (decitabine) and 92% CR/CRi in Arm B (control). The patient who discontinued study participation on day 6 after receiving all decitabine doses and only one dose of cytarabine remained in a complete remission for two months without any further leukemia-directed treatment. She eventually resumed standard therapy two months later and remains in CR 56+ months later. MRD by multi-parameter flow cytometry showed that 85% of patients in Arm A were MRD negative at day 30 of induction compared to 67% in Arm B. In a follow-up of patients as of August 2015, 8 of 14 (57%) in Arm B and 5 of 10 (50%) of evaluable patients in Arm A relapsed following additional treatment.

### Biological marker analysis

Quantitative DNA methylation analyses revealed global changes in methylation following decitabine priming. Diagnostic and end-induction marrows were analyzed in 9 patients in each Arm (**Figure S1**). Paired differential methylation analysis of end-induction marrows to patient matched screening marrows revealed 6990 differentially methylated CpG loci (DML) encompassing 2518 genes in Arm A compared to only 1090 DML (539 genes) in Arm B (**Tables S1A-B**). Only DML in Arm A (n=4597) survived false discovery p-value correction. Of all DML in Arm A, 4134 were hypomethylated and 2856 were hypermethylated. In Arm B, 785 DML were hypomethylated and 305 were hypermethylated. There were 795 DML (438 genes) common to both arms. Although about 80% of genes altered by DNA methylation in Arm B were common to Arm A, there were significantly more probes altered for a given gene in Arm A. Moreover, 78% of hypomethylated probes in Arm B were common with Arm A, compared with 56% of hypermethylated probes common between treatment arms. The median delta-beta values for Arms A and B were -0.27 and -0.28, respectively, indicating modest overall hypomethylation induced by either treatment regimen at the specified delta-beta cut-off. Forty-one percent of DML were hypermethylated after decitabine therapy compared with 28% after chemotherapy only. Regional and functional CpG distribution of DML after therapy in both treatment arms was also examined. Functional distribution relates CpG position to transcription start sites (TSS) −200 to −1500 bp, 5’ untranslated region (UTR), and exon 1 for coding genes as well as gene bodies. In both treatment arms, gene body hypermethylation was the most frequent change, followed by gene body hypomethylation and TSS 200 hypomethylation (**Figure S2**). Regional distribution of DML was assessed based on proximity to the closest CpG island. In addition to CpG islands, shores are 0-2 kb from CpG islands, shelves are 2-4 kb away, and open sea regions are isolated loci without a designation. CpG island hypomethylation occurred in greater than 68% of DML in both groups. Hypermethylation occurred most prominently in open sea regions and to a greater degree in Arm A patients compared with those in Arm B receiving chemotherapy alone (**Figure S3**).

Unsupervised clustering analysis of DML for both treatment arms demonstrated strong separation of screening and end-induction marrows except for one sample pair in Arm A and three sample pairs in Arm B (**Figure 2**, top panels). In two of these cases, pre- and post-treatment samples co-clustered with its matching sample. One case in Arm A and Arm B clustered with diagnostic marrows, suggesting the marrow was possibly unaffected by therapy and indeed the sample in Arm A (1006_1004) was from a patient with stable disease. Overall, these data indicate that decitabine therapy has a homogenizing effect on the recovering end-induction marrow in AML. This was evident when compared with Arm B samples, where DNA methylation was more heterogeneous after standard chemotherapy treatment.

**Figure 2.**
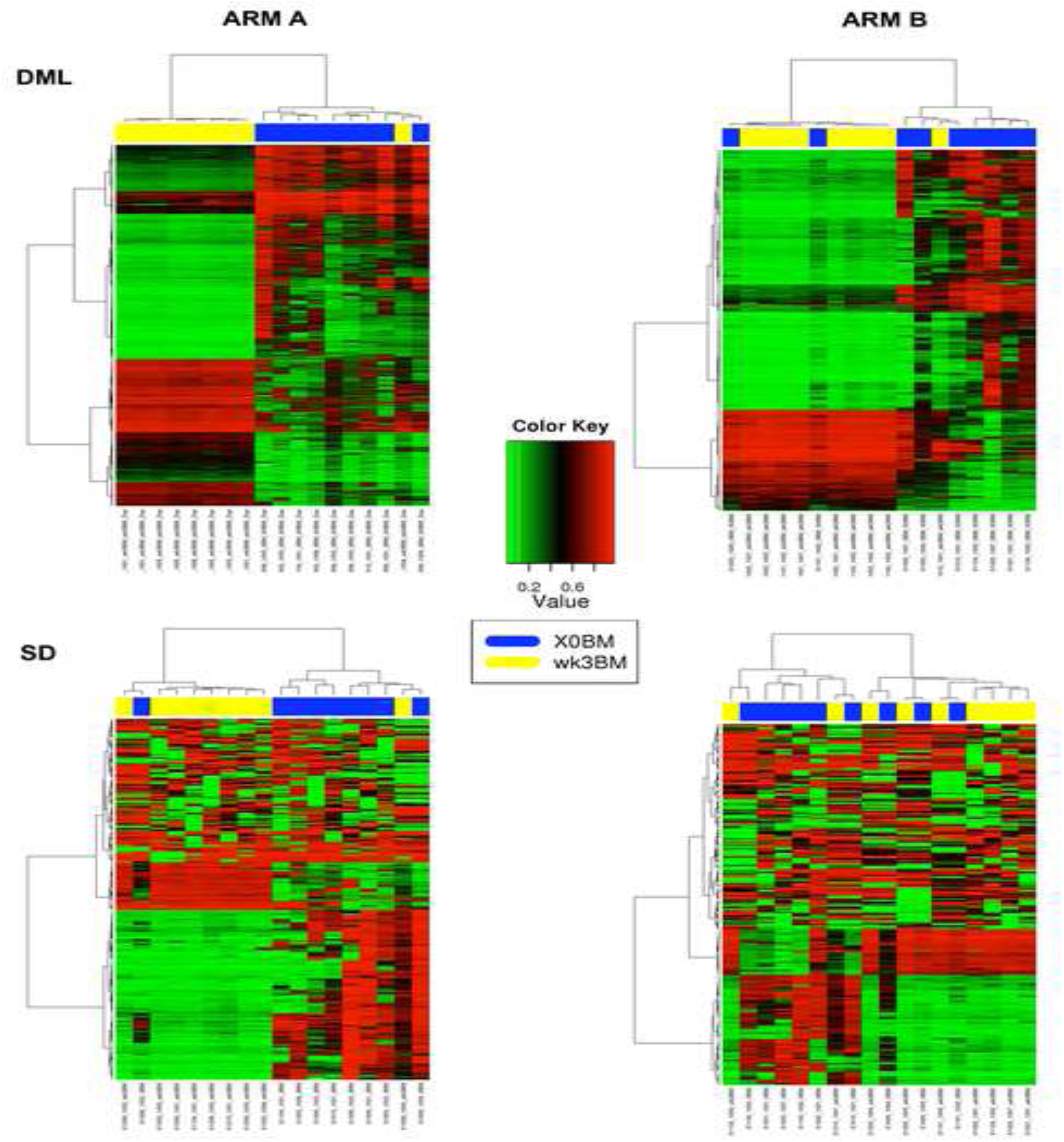
Hierarchical clustering of differentially methylated loci (DML) in Arm A (decitabine + chemotherapy) and Arm B (chemotherapy alone). Top Panel: unsupervised clustering analysis of 6990 DML in Arm A and 1090 DML in Arm B revealed separation of end of induction recovering marrows at week 3 from screening marrows (X0BM). Lower Panel: Unsupervised hierarchical clustering of the top 0.1% most variable loci (SD) also separated screening marrows (X0BM) from end of induction recovery marrows at week 3. In both analyses, Arm A shows better separation of the two time points than in Arm B, indicating that decitabine therapy appears to homogenize marrows to a larger degree than in chemotherapy alone.

To further assess the changes in recovering marrows in both arms, we performed an unsupervised clustering analysis of the top 0.1% most variable CpG probes (~430 probes) in the post-processed data. These data confirmed that the end-induction recovering marrows were distinctive from screening marrows and more homogenized in the decitabine-treated arm compared with those in the control arm (**Figure 2**, lower panels). To ensure that samples were not molecularly different at screening between the arms, we performed the above analyses comparing screening marrows in Arm A with Arm B and observed only 492 DML. Of these, 291 were common to the Arm A DML list, while 3 DML were common to both Arm A and B comparisons and 0 DML were common to Arm B DML (**Figure 3**). These intrinsically different loci were excluded from downstream gene-level analysis. Among the genes most prevalently hypomethylated in Arm A were FOXG1, VSTM2A, WT1, ZNF135, ZIC1, and ZIC4, (**Figure 4**), which may potentially be used to measure decitabine activity. In addition, time-dependent promoter hypomethylation of these genes also occurred in peripheral blood lymphocytes (**Figure 5**), confirming their significance as potential biomarkers of decitabine response. Most notably, recovery of promoter methylation in peripheral blood was seen in a patient with stable disease and whose recovering marrow co-clustered with diagnostic marrow, hinting at signs of preliminary efficacy. The data point to the potential utility of these genes as biomarkers of minimal residual disease in patients treated with decitabine.

**Figure 3.**
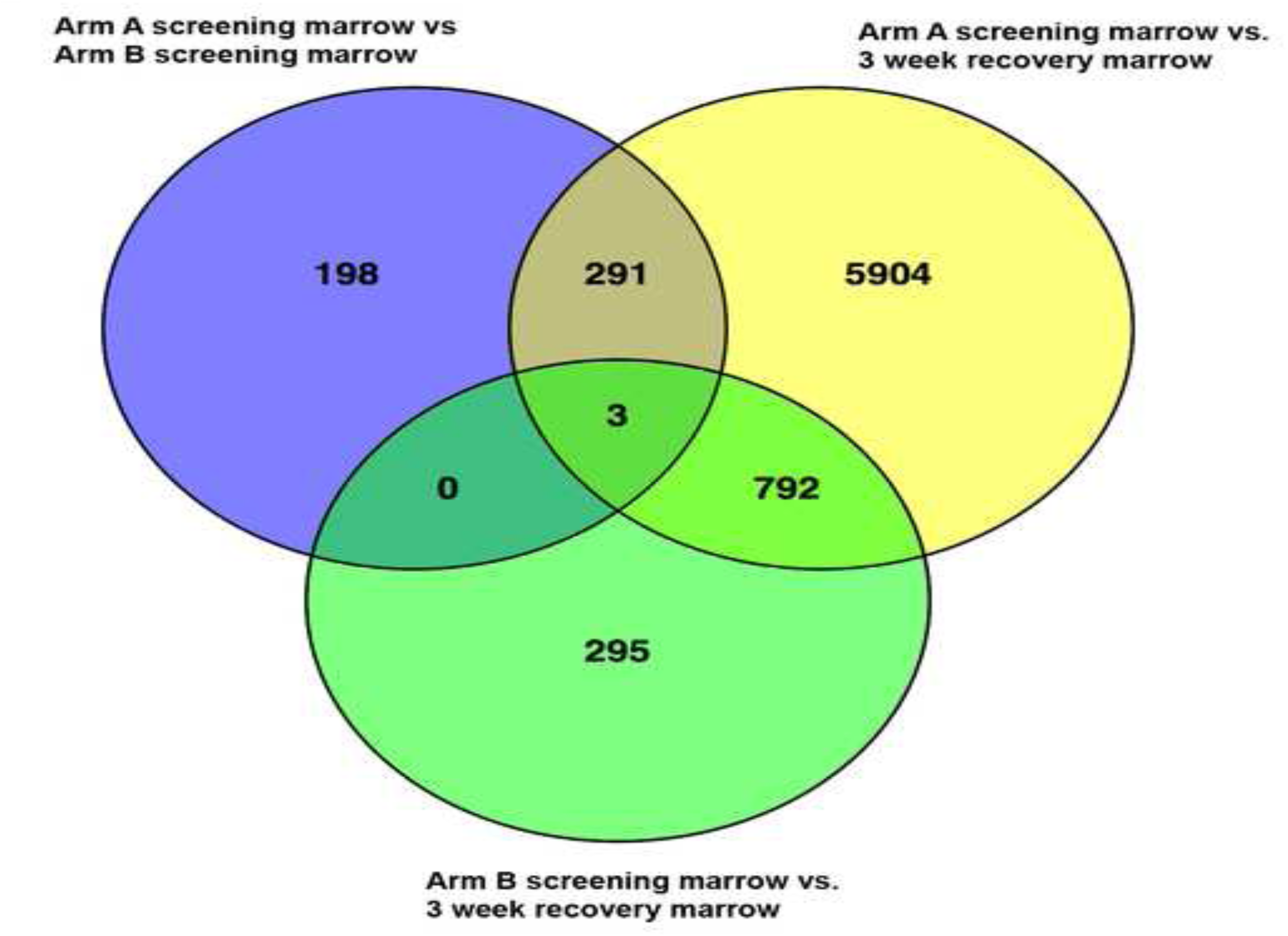
Overlap of differentially methylated loci between arms and time points in Arm A (DADE), Arm B (ADE), and screening vs. recovery marrow aspirates. Screening marrows for samples in Arm A and Arm B are also compared and demonstrate little intrinsic bias between groups.

**Figure 4.**
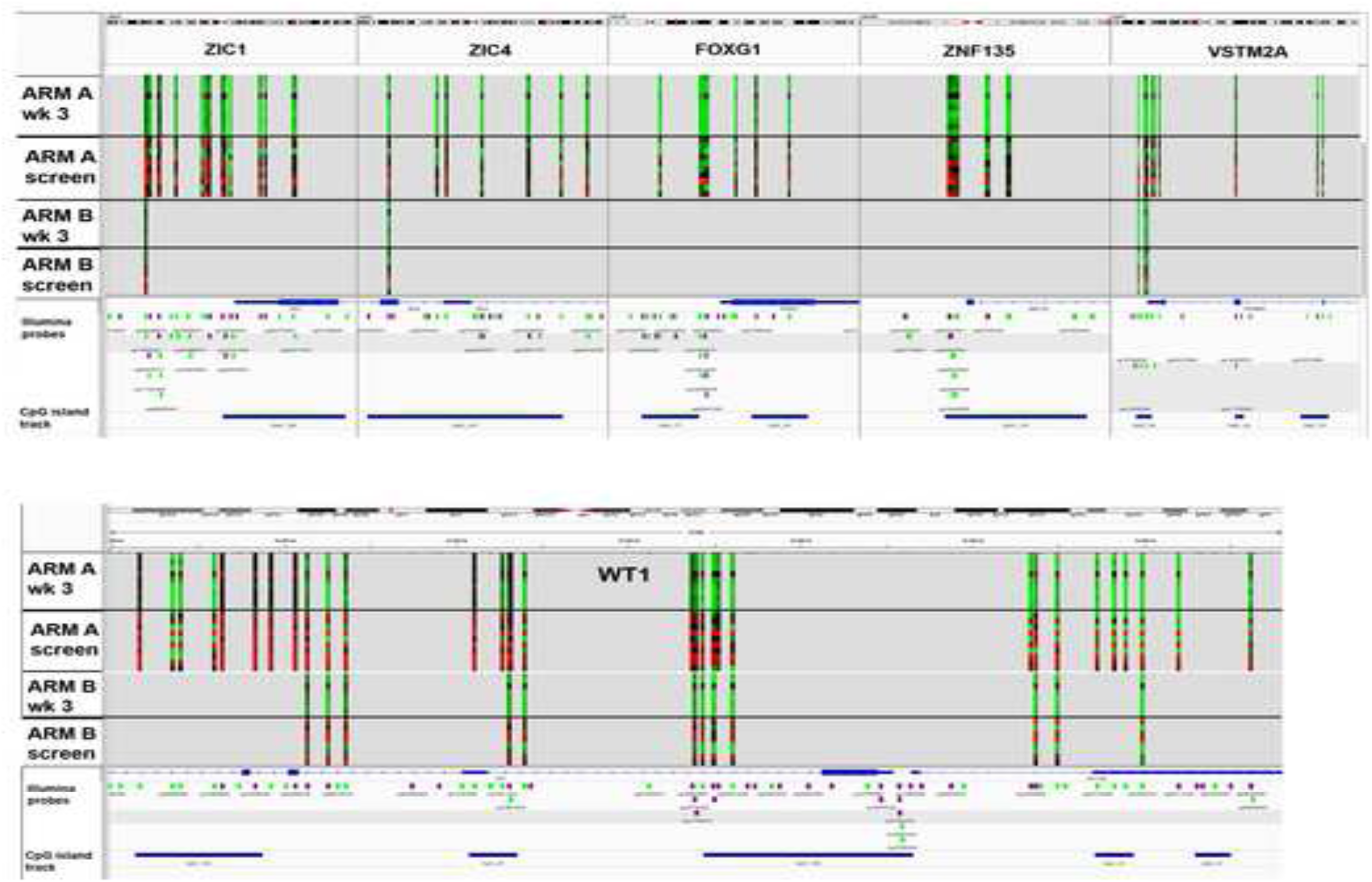
Integrated Genomic Viewer snapshot of differentially methylated genes affected by hypomethylation in response to decitabine therapy. Vertical heatmaps represent significantly differentially methylated (p value <0.05) probes in the 6 genes illustrated. Each row on the heatmap represent a unique sample. Many more probes were differentially methylated in Arm A (decitabine + chemotherapy) compared with Arm B (chemotherapy alone) for the probes shown. Hypomethylation (green) in response to decitabine (Arm A) is evident in end of induction recovery marrows (wk 3) compared with diagnostic marrows (screen). Array avg β values are represented in the heatmap. Scale ranges from 0-1, where 0 is unmethylated (green) and 1 is fully methylated (red). Tracks shown: gene, CpG 450K probe, and CpG island.

**Figure 5.**
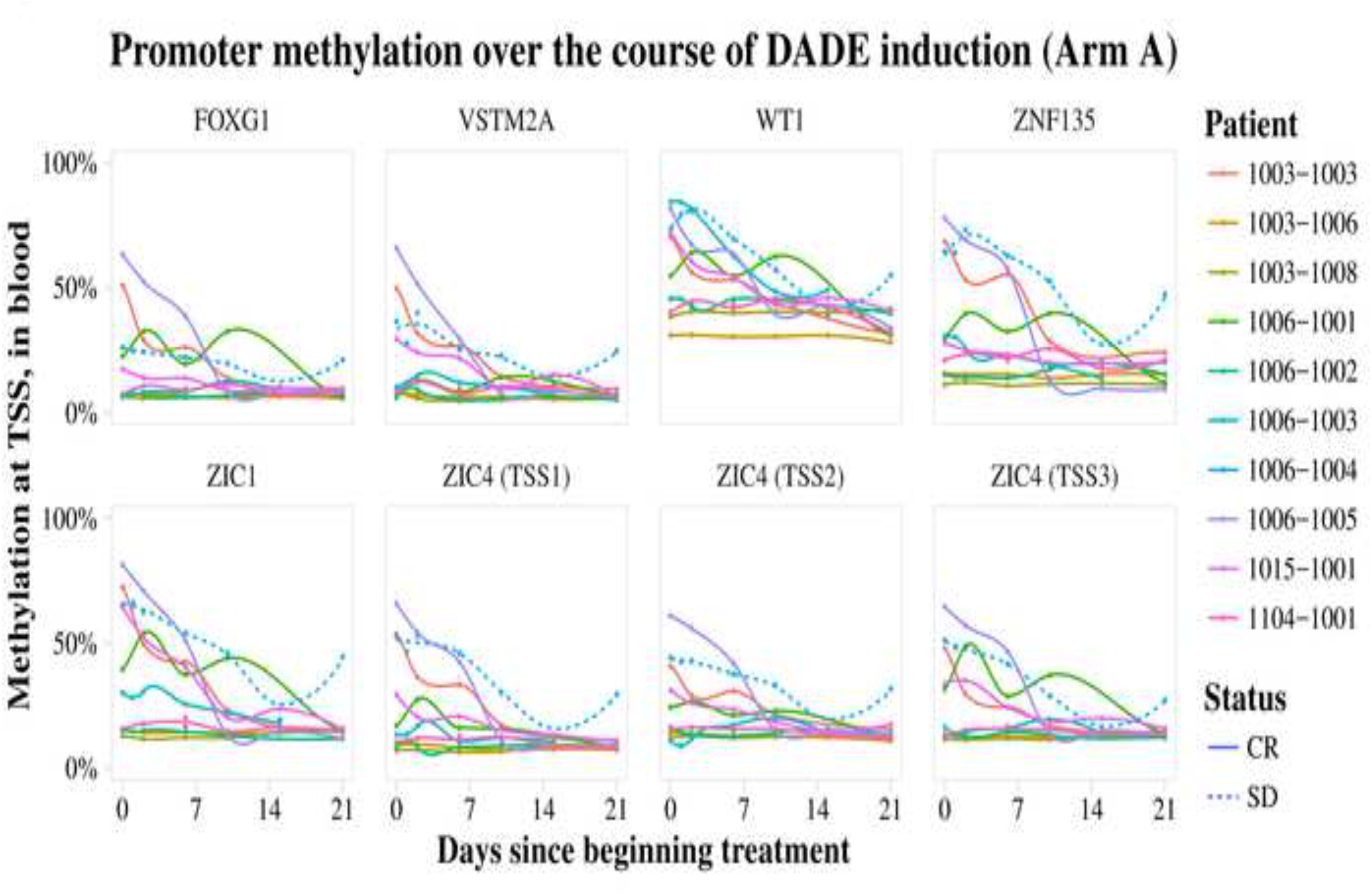
Time-course collection of peripheral blood samples in Arm A reveal consistent decreases in promoter methylation at relevant transcription start sites over treatment in all responders, as well as a reversal of this decrease in the sole non-responsive patient (1006-1004). A distinct uptick in the patient’s promoter methylation from day 14 to 21 is noted, which corresponded clinically to the patient’s disease progression.

**Table 4** shows DNA methylation changes in several key biological pathways potentially important to response to decitabine and chemotherapy. The top canonical pathways in IPA for Arm A DML included gene alterations affecting mostly neuronal signaling such as neuropathic pain signaling and glutamate receptor signaling (**Table 4**). In Arm B, the top IPA canonical pathways included DML affecting embryonic stem cell signaling and Rho GTPase signaling (**Table 4**).

**Table 4.** Ingenuity Pathway Analysis of Differentially Methylated Genes.

|  | p-value | Overlap n (%) |
| --- | --- | --- |
| **Arm A Top Canonical Pathways** |  |  |
| Neuropathic Pain Signaling in Doral Horn Neurons | 7.5e-12 | 35/100 (35.0%) |
| Glutamate Receptor Signaling | 2.45e-11 | 25/57 (43.9%) |
| Amyotrophic Lateral Sclerosis Signaling | 2.05e-09 | 31/98 (31.6%) |
| Synaptic Long Term Potentiation | 1.00e-06 | 30/119 (25.2%) |
| Hepatic Fibrosis/Hepatic Stellate Cell Activation | 1.01e-06 | 40/183 (21.9%) |

| Arm B Top Canonical Pathways |  |  |
| --- | --- | --- |
| Transcriptional Regulatory Network in Embryonic Stem Cells | 2.68e-04 | 4/40 (10.0%) |
| Thrombin Signaling | 7.94e-04 | 7/191 (3.7%) |
| GPCR-Mediated Integration of Enteroendocrine Signaling Exemplified by an L Cell | 2.36e-03 | 4/71 (5.6%) |
| Signaling by Rho Family GTPases | 2.54e-03 | 7/234 (3.0%) |
| CXCR4 Signaling | 7.03e-03 | 5/152 (3.3%) |

Differentially methylated genes for Arm A (decitabine + chemotherapy; n=2518) and Arm B (chemotherapy alone; n=539) were entered into the core pathway analysis option. Top 5 canonical pathways are shown along with a significant p-value and the number of genes in each list belonging to the pathway.

This study is unique in comparing the epigenetic and RNA expression consequences of induction chemotherapy following pre-treatment with decitabine vs chemotherapy alone in end-induction recovering bone marrow. This comparison revealed 104 differentially expressed genes in Arm A (**Table S2A**) and 74 genes in Arm B (**Table S2B**). There are 5 genes common to both treatment arms. Twelve genes that were differentially expressed in Arm A had DNA methylation changes affecting 72 DML (**Table S3A**,). Among these were 57 probes impacting PCDGHGB2, the potential calcium-dependent cell-adhesion protein; the gene was hypomethylated but downregulated. In Arm B there were only 23 DML (**Table S3B**) corresponding to eight differentially expressed genes. Only *SIX3* overlapped between both arms at a single DML. Additionally, we saw no evidence of sustained repeat element reactivation in recovering week three marrows with or without decitabine pre-treatment (**Figure 6**).

**Figure 6.**
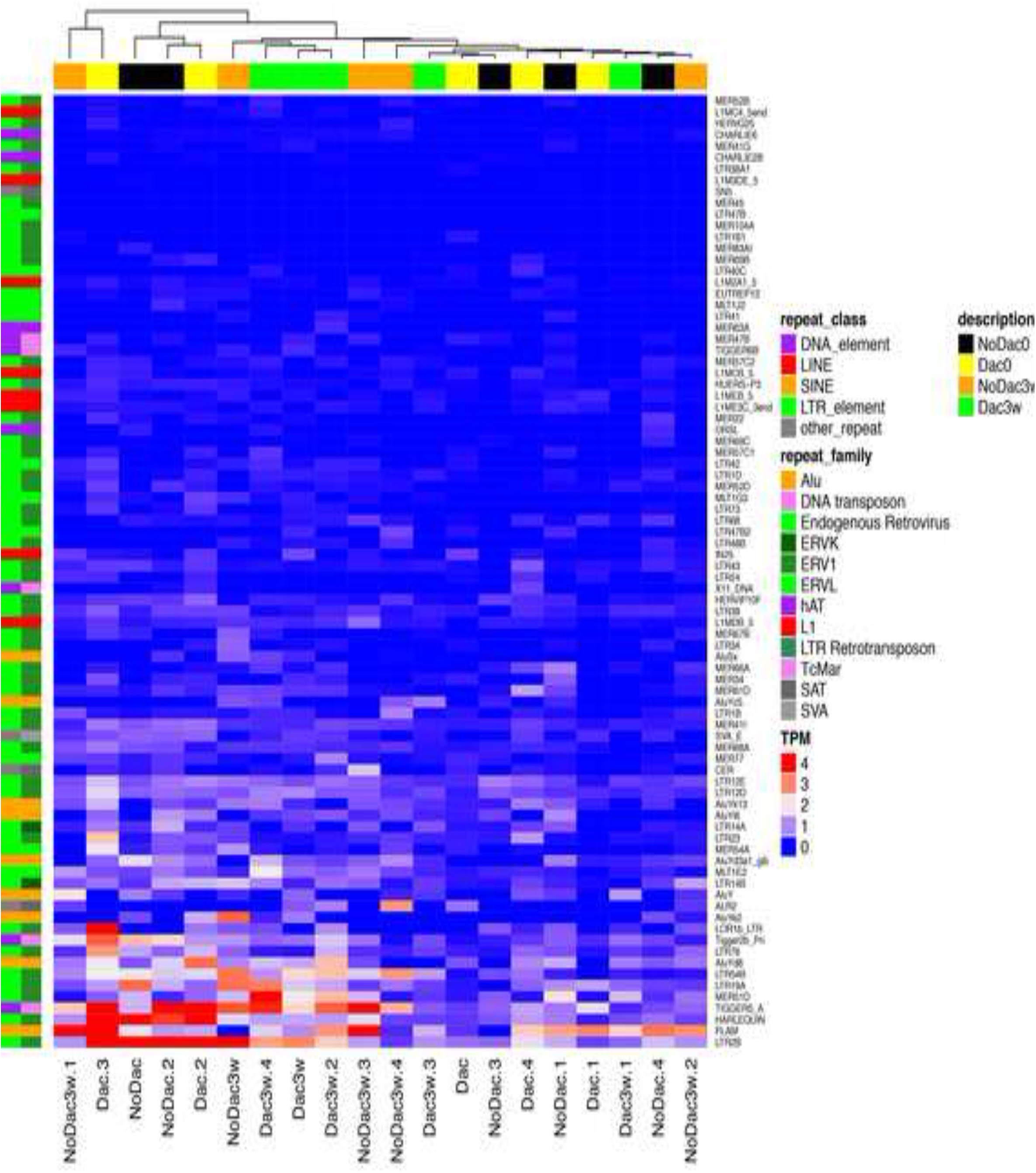
Unsupervised hierarchical clustering of repeat element expression in screening and week 3 bone marrows, with (Arm A) or without (Arm B) decitabine priming, in transcripts per million (TPM). No clear separation by trial arm is visible, and a maximum of 4 transcripts per million is observed, suggesting that mobilization of repeats by demethylating agents does not persist beyond the course of treatment and is not significantly different between the treatment arms.

## DISCUSSION

This first-in-pediatric randomized trial of epigenetic priming in children with newly diagnosed AML demonstrated safety and tolerability and establishes the feasibility required to develop future trials for assessing durability of response. We did not assess durable responses in this study because its main purpose was to first assess safety and tolerability of decitabine in children. The results trended towards non-inferiority of morphologic, immunophenotypic, and MRD response, although the small sample size limited statistical analyses. There was also evidence of decitabine-induced effects in end-induction bone marrow aspirates compared with those obtained at diagnosis. Children treated with decitabine as a single agent for five days prior to conventional cytotoxic therapy did not have rapid progression of their leukemic burden during the pre-phase, further supporting the feasibility and safety of this approach. Based on these results, a dose of 20 mg/m^2^ daily for 5 days prior to standard conventional chemotherapy could be considered for testing for durable responses.

Most non-hematologic AEs reported in this study were mild to moderate in severity (grade 1 or 2), reversible, and easily managed. In general, the safety profile of decitabine in children with AML was consistent with that seen in adults. No previously unreported decitabine toxicities were observed. Drug-related hematologic toxicity, anorexia, and asymptomatic grade 3 hypokalemia and hypophosphatemia were slightly more common in decitabine treated patients.

This trial was not powered to detect a difference in response between the two arms and treatment with decitabine prior to standard induction therapy resulted in a similar morphological response compared to standard induction therapy. Of note, there were more high-risk cytogenetic patients in the decitabine arm (4 versus 0), which may indicate a benefit for decitabine priming prior to treatment in these patients. Patients in this study were generally representative of children with *de novo* childhood AML in age, gender, and biologic features, however there were slightly more patients with *WT1* and *CEBP*A mutations than previously reported. In addition, the impact on MRD suggests that decitabine may induce a higher percentage of MRD negative remissions; however, again the numbers in this study are too small to calculate statistics in a meaningful way. There was a non-significant trend toward a longer time to recover neutrophil and platelet counts in the decitabine-treated patients (data on file), and no AEs or SAEs were noted as a result of delays in count recovery. These results suggest that exposure to decitabine may have important implications for sensitizing potentially resistant leukemic clones to cytotoxic chemotherapy, resulting in deeper remissions predictive of more favorable outcomes. Larger randomized studies are needed to confirm these findings.

Alterations in DNA methylation and RNA expression patterns in end-induction bone marrow between the two groups of patients suggest important consequences of exposure to decitabine pretreatment on hematopoietic recovery after exposure to intensive chemotherapy, where decitabine appears to have a homogenizing effect on end-induction marrows that may ultimately be important in priming for chemotherapy sensitization. Our data suggest that while chemotherapy alone may have an effect on DNA methylation, the effect is clearly augmented by the addition of decitabine via epigenetic changes that may impact both leukemic, normal hematopoietic progenitors, as well as bone-marrow stromal cells. Furthermore, the data suggest that decitabine therapy can be used to measure MRD/or patient response to therapy by assessing DNA methylation status of specific promoter regions non-invasively in blood. The pathway analysis conducted to compare differentially methylated loci in the decitabine treated arm to the chemotherapy only arm revealed a number of pathways implicated to neuronal signaling but the results were quite distinct from the ADE only arm. While the implications of neuronal signaling are not currently clear, we postulate it must be related to the bone marrow niche post-treatment since no dorsal root ganglia or other neuronal tissue exist in the marrow. Presumably this could be due to ion channel currents that play an important role in bone-marrow derived mesenchymal stem cells and hematopoietic progenitors (26). Our observations further suggest that monitoring changes in normal progenitors may be important in understanding the short and longer-term consequences of exposure to methyltransferase inhibitors on malignant and normal bone marrow progenitors.

An increased percentage of DMLs were hypermethylated in patients receiving decitabine compared to those receiving chemotherapy alone, suggesting that decitabine has an effect on the methylome of the recovering marrow beyond DNA demethylation. This appears to result in changes that differ both qualitatively and quantitatively from chemotherapy alone. Importantly, chemotherapy molecularly alters the recovering marrow epigenome, but the number of changes is significantly fewer than with decitabine and the pathways affected differ significantly.

## CONCLUSIONS

The toxicity and PK results observed in the patients in this study suggest that decitabine can be safely combined with standard doses and schedules of anticancer agents in children with newly diagnosed AML. Furthermore, our data suggest that this regimen alters DNA methylation and RNA expression compared to ADE chemotherapy alone, and patients treated with decitabine could have minimal residual disease measured by assessing DNA methylation status of specific promoter regions. Preclinical studies have shown additive or synergistic activity when decitabine is combined with a variety of other anticancer therapies (27-30) and results from trials such as this provide further evidence of feasibility, safety and possible strategies for larger randomized trials in patients with newly diagnosed or recurrent/refractory leukemia as well as in the minimal disease state during post-remission follow-up. No excess or unexpected toxicities were seen. The most common drug-related grade 3 or grade 4 AEs were hematologic and PK/PD were as expected. Complete remission rates were similar. Patients treated with decitabine prior to conventional chemotherapy had distinct changes in DNA methylation and transcriptional regulation. Repeat element transcription may be of interest for further mechanistic study. In conclusion, epigenetic therapy with decitabine is safe for use in children and the clinical findings together with molecular correlative studies suggest that there may be early signs of enhanced efficacy. However, further studies are needed to definitively determine the long-term patient outcomes of decitabine priming in children with AML.

## LIST OF ABBREVIATIONS

ADE = cytarabine/daunorubicin/etoposide chemotherapy regimen

AML = acute myelogenous leukemia

AUC = area under the curve

BSA = body surface area

C max = maximum plasma concentration

CML = chronic myelogenous leukemia

CNS = central nervous system

CR = complete remission

CR = complete remission with incomplete count recovery

DADE = decitabine plus daunorubicin/cytarabine/etoposide chemotherapy regimen

DML = differentially methylated loci

DNA = deoxyribonucleic acid

FAB = French-American-British classification

HIV = human immunodeficiency virus

LFS = leukemia-free survival

MRD = minimal residual disease

OS = overall survival

PD = pharmacodynamics

PK = pharmacokinetics

RNA = ribonucleic acid

Tmax = time to maximum plasma concentration

## DECLARATIONS

### Ethics Approval and Consent to Participate

The study protocol was approved by each institutional review boards at every participating site and was conducted in accordance with the Declaration of Helsinki, Good Clinical Practice, and all local and federal regulatory guidelines. A parent or legal guardian provided written informed consent, with patient assent as appropriate according to institutional requirements.

### Consent for Publication

Not applicable

### Availability of Data and Materials

The datasets generated and analyzed in this study are included in this article and are available from the corresponding author on reasonable request. In addition, the DNA methylation and RNA-seq data discussed in this publication have been deposited in NCBI’s Gene Expression Omnibus, and are accessible through the GEO series accession number GSE78963.

### Authorship Contributions

L.G. and R.J.A. designed and wrote the study concept and scientific hypotheses, derived the full protocol, provided study oversight during the conduct of the study, enrolled patients on the study, reviewed data and toxicity of patients, and interpreted the data. R.J.A. performed many of the scientific experiments and contributed to a draft of the manuscript. L.G. performed the literature search, wrote the first, all drafts and the final manuscript, helped prepare and reviewed the Figures and Tables, and collated and incorporated comments from all co-authors. B.S. conducted experiments and provided oversight of the scientific experiments not completed by R.J.A., prepared and reviewed the Figures and Tables, and contributed substantially to the data analysis and writing and editing of the manuscript. T.T. helped analyze and review the experiments and Figures and Tables, and assisted substantially in the writing and reviewing of the final manuscript. J.E.F., D.W., C.L., G.C.G., W.S.L., J.C., D.L., and S.M. contributed to the scientific experiments and data analyses including Figure preparation and review. J.E.F. and T.T. also contributed to writing the Methods sections, reviewing, and editing the manuscript. S.M. contributed to the data analysis and commentary. F.A., M.E.M., C.A., P.B., T.C., L.M., A.N., J.P., J. B. enrolled patients on the clinical trial and reviewed and commented on the manuscript prior to submission. M.J., L.C., S.S., and P.T. reviewed the study and manuscript. M.J. representing the Eisai Inc., provided study enrollment materials, oversight and analysis within the context of the protocol operations and data analysis. Eisai, Inc., is the manufacturer of decitabine and the company providing partial support for the conduct of this trial.

### Dedication

The authors of this paper would like to recognize the fundamental contributions that Dr. Robert Arceci made not only in the conception, design, and conduct of this study and the associated biology work incorporated within, but to the field of pediatric oncology as a whole. During the laboratory correlate analysis and preparation of this manuscript, Dr. Arceci’s life was tragically cut short in a fatal traffic accident. His collaborators on this study wish to acknowledge his myriad contributions to this work and the field, and remain dedicated to pursuing the highest degree of scientific collaboration and integrity in his honor.

## Acknowledgements

Dr. Francoise Mechinaud (Royal Children’s Hospital, Australia) and Dr. Wendy Tcheng (Children’s Hospital of Central California) also treated patients on this trial. Dr. Peter Tarrasoff had oversight and analysis of the protocol operations on behalf of Eisai, Inc., the manufacturer of decitabine and the company providing partial support for the conduct of this trial.

Lori Cuyugan and Shobana Sekar (Translational Genomics Research Institute, Phoenix, AZ) for the generation and analysis of RNA-seq data.

## SUPPLEMENTARY FIGURE LEGENDS

**Figure S1.** Schema of sample analysis workflow.

**Figure S2.** Distribution of differentially methylated loci (DML) according to functional CpG contextual distribution in Arms A (decitabine + chemotherapy) and B (chemotherapy alone). Pie charts demonstrate the frequency by which hyper or hypomethylated loci are distributed according to their functional position.

**Figure S3.** Distribution of differentially methylated loci (DML) according to CpG Island contextual distribution in Arms A (decitabine + chemotherapy) and B (chemotherapy alone). Pie charts demonstrate the frequency by which hyper or hypomethylated loci are distributed according to their proximity of CpG islands.

